# COVID-19 vaccine booster induces a strong CD8^+^ T cell response against Omicron variant epitopes in HLA-A*02:01^+^ individuals

**DOI:** 10.1101/2022.01.12.473243

**Authors:** Andrea T. Nguyen, Christopher Szeto, Demetra S.M. Chatzileontiadou, Zhen Wei Marcus Tong, Michael J. Dewar-Oldis, Lucy Cooper, Lawton D. Murdolo, Keng Yih Chew, Katie E. Lineburg, Alan Riboldi-Tunicliffe, Rachel Williamson, Bradley J. Gardiner, Dhilshan Jayasinghe, Christian A. Lobos, You Min Ahn, Emma J. Grant, Corey Smith, James McMahon, Kim L. Good-Jacobson, Peter J. Barnard, Kirsty R. Short, Stephanie Gras

## Abstract

The >30 mutated residues in the Omicron spike protein have led to its rapid classification as a new SARS-CoV-2 variant of concern. As a result, Omicron may escape from the immune system, decreasing the protection provided by COVID-19 vaccines. Preliminary data shows a weaker neutralizing antibody response to Omicron compared to the ancestral SARS-CoV-2 virus, which can be increased after a booster vaccine. Here, we report that CD8^+^ T cells can recognize Omicron variant epitopes presented by HLA-A*02:01 in both COVID-19 recovered and vaccinated individuals, even 6 months after infection or vaccination. Additionally, the T cell response was stronger for Omicron variant epitopes after the vaccine booster. Altogether, T cells can recognize Omicron variants, especially in vaccinated individuals after the vaccine booster.

**One-Sentence Summary:** CD8^+^ T cells response against Omicron variant epitopes is stronger after the vaccine booster.

---

The new SARS-CoV-2 variant named Omicron (B.1.1.529) was first isolated in Southern Africa, and rapidly classified as a variant of concern (VOC) by the WHO (*1, 2*). Omicron has >30 spike mutations, 21 of which are not shared with the other classified variants (GISAID (*3*)). The first reports on the ability of neutralizing antibodies (nAb) to block the Omicron variant were alarming. The data showed a lack of neutralization in both vaccinated (*4*) and recovered individuals (*5, 6*); a 40-fold decrease of nAb levels against Omicron in vaccinees (*5, 7*). The study by Sigal’s team showed that vaccinated individuals with prior SARS-CoV-2 infection had higher levels of nAb to Omicron than individuals who were vaccinated alone (*5*). In addition, a booster vaccine (3^rd^ vaccine dose) stimulated cross-reactive nAb, limiting the nAb levels decrease to a 4-6 fold compared to the original SARS-CoV-2 (*8*). Overall, the current data suggests a decreased nAb levels against Omicron, with a serious potential for viral escape from this variant.

It is clear that nAb offers protection against SARS-CoV-2, however the immune system is composed of multiple immune cell subsets with the potential to both recognize and provide protection against emerging variant strains. T and B lymphocytes have important roles in antiviral immunity, including against SARS-CoV-2, with memory capabilities that offer long-lasting immunity (*9–13*). T cells in SARS-CoV-1 recovered individuals demonstrated retention of a robust cross-reactive memory response persisting after 17 years (*13*). Cross-reactive T cells have also been identified in COVID-19 recovered and unexposed individuals (*14, 15*), suggesting that memory T cells could provide long lasting protection. Meanwhile, despite waning SARS-CoV-2-specific antibody titers, studies demonstrate that memory B cells have a long lasting presence after infection and vaccination (*10, 16*), and exhibit cross-reactive potential to varying degrees (*17*). Overall, both T and B cells could be essential in cross-reactive recognition of the Omicron variant.

A global effort is being made to understand the immune response to SARS-CoV-2 in depth, mediated by vaccination or infection, with details that would give researchers an advantage against future variants. Here, we show that, in HLA-A*02:01^+^ individuals, the nAb levels against the original SARS-CoV-2 virus increase after the 2^nd^ dose of vaccine (ChAdOx1 nCoV-19 [AstraZeneca] or the BNT162b2 [Pfizer] vaccine) then slowly decrease over 6 months after infection or vaccination, with a similar pattern for memory B cells. Our study focuses on spike-derived CD8^+^ T cell epitopes restricted to HLA-A*02:01, one of the most common Human Leukocyte Antigen (HLA) class I molecules (15.3% frequency worldwide and ~40% in Caucasian populations (*18*)). Five of the Omicron mutations in the spike protein fall within three HLA-A*02:01-restricted peptides, which could escape CD8^+^ T cell recognition. To understand the impact of Omicron mutations we solved the crystal structures of peptide-HLA-A*02:01 complexes (pHLA-A*02:01) and assessed CD8^+^ T cell recognition to the three Omicron variants in recovered and vaccinated individuals.

We firstly assessed the level of spike-specific nAb, against the original SARS-CoV-2 virus, in COVID-19 recovered HLA-A*02:01^+^ individuals early (< 2 months) and late (up to 8 months) after SARS-CoV-2 infection (**Table S1**). While we observed a decrease of the median value over time (**Fig 1A**), the nAb were retained, in line with other studies (*19, 20*). We also determined the nAb levels in individuals after vaccination (**Table S1**) with samples isolated from blood samples collected 10 to 14 days after the 1^st^ and 2^nd^ dose, and 6 months post-1^st^ dose of the vaccine. The nAb levels were significantly increased after the 2^nd^ dose of either vaccine (*p* = 0.0039) and decreased, but still present, after 6 months to comparable levels post-1^st^ vaccine dose (**Fig 1B**), in line with previous reports (*21*). As the nAb levels decreased over time (*21, 22*), we wanted to know if the memory B cell pool would be maintained over time, helping provide long term protection. The level of spike-specific memory B cells was determined in COVID-19 recovered individuals (**Fig 1C** and **Fig S1**), and vaccinated individuals (**Fig 1D** and **Fig S1**) at different time points. The level of spike-specific memory B cells only slowly decreased over time in both recovered and vaccinated individuals, as previously shown (*10, 21*), with a noticeable increase in memory B cells after the 2^nd^ vaccine dose (**Fig 1C-D**).

**Fig. 1.**
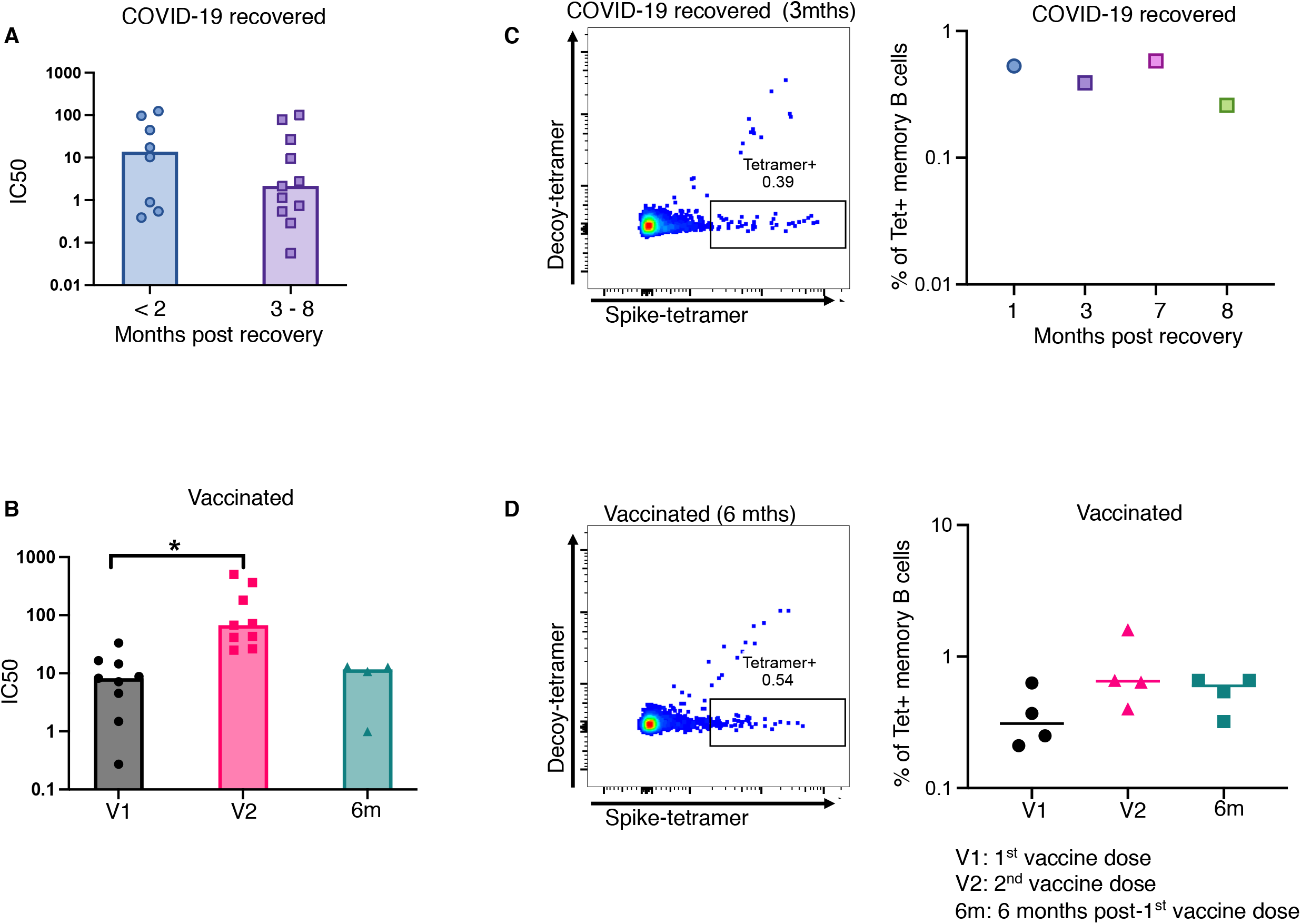
Antibody and B cell responses towards spike protein derived from the original SARS-CoV-2 in recovered and vaccinated individuals. **A.** Plasma reactivity to spike protein in COVID-19 recovered individuals at <2 months and 3-8 months post infection time points. **B.** Plasma reactivity to S protein in vaccinated individuals at three post vaccination time points (1^st^ and 2^nd^ vaccine dose and 6 months post-1^st^ vaccine dose). **C.** Spike tetramer positive B cells in COVID-19 recovered individuals at different time points post recovery. **D.** Spike tetramer positive B cells in vaccinated individuals at three post vaccination time points. The decoy tetramer used on panels (**C-D**) was of HLA-A*02:01-M158 (M158 is an influenza epitope).

Given that preliminary studies showed a partial escape of Omicron from nAb (*4–6, 8*), we wanted to determine the impact of Omicron spike mutations on the T cell response. Three spike peptides able to bind HLA-A*02:01 are mutated in Omicron (**Table 1**). The first two peptides were described as immunogenic from the original SARS-CoV-2, the S417 (*23, 24*) and S976 (*25, 26*), and have a single mutated residue in Omicron, with the S417 variant also shared in the Beta variant. The third peptide, S367, we predicted to bind HLA-A*02:01, has three Serine residues mutated to Leu, Pro and Phe at positions 5, 7, and 9, respectively (**Table 1** and **Fig S2**).

**Table 1.**
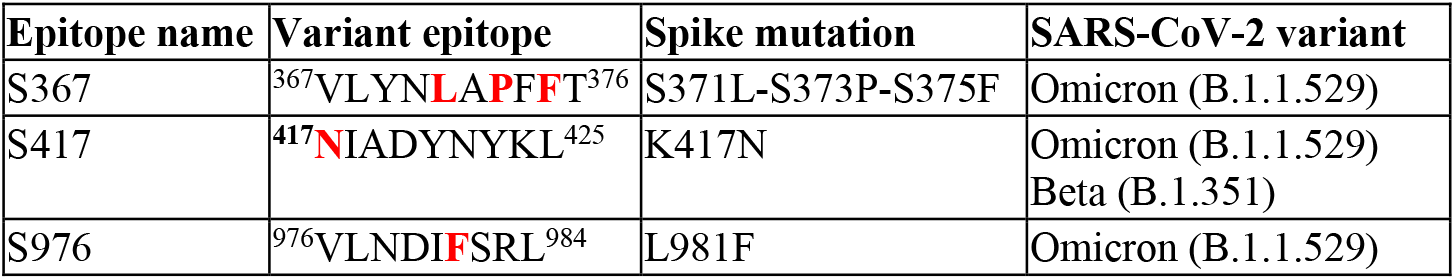
Spike derived HLA-A2-restricted epitopes from the Omicron variant. The mutated residues present in the Omicron variant of SARS-CoV-2 are indicated in bold and red.

| **Epitope name** | **Variant epitope** | **Spike mutation** | **SARS-CoV-2 variant** |
| --- | --- | --- | --- |
| S367 | <sup>367</sup>VLYN**LAPFFT**<sup>376</sup> | S371L-S373P-S375F | Omicron (B.1.1.529) |
| S417 | <sup>417</sup>**N**IADYNYKL<sup>425</sup> | K417N | Omicron (B.1.1.529)<br>Beta (B.1.351) |
| S976 | <sup>976</sup>VLNDI**F**SRL<sup>984</sup> | L981F | Omicron (B.1.1.529) |

To understand the impact of these Omicron spike mutations on peptide presentation by HLA-A*02:01, we solved the structures of S367, S976 and S417-Omicron (S417-O) bound to HLA-A*02:01 (HLA-A*02:01-S417 was previously solved (*24*)), and generated models for S367-Omicron (S367-O) and S976-Omicron (S976-O) (**Fig 2**, **Table S2**, and **Fig S3**). The overall structure of the HLA-A*02:01 cleft (residue 1-180) bound to S367, S976 and S417-O was similar with an average root mean square deviation (r.m.s.d.) of 0.14 Å. The three peptides bind in the cleft in a similar conformation to other HLA-A*02:01-restricted peptides (*27, 28*). As predicted the S367 peptide could bind to HLA-A*02:01, however, there is currently no data on the S367 peptide. The S367 peptide shows that P5-S and P9-S are solvent exposed, and as such accessible for TCR interaction, whereas P7-S is buried in the cleft forming a hydrogen bond with Arg97 (**Fig 2A** and **Fig S4A**). Based on the structure of S367, a model of S367-O was generated by mutating the three serine residues to P5-L, P7-P, and P9-P (**Fig 2B**). While these substitutions did not introduce steric clashes, the addition of a P7-P could alter the S367-O peptide conformation. Interestingly, the triple mutations significantly increased the stability of the HLA-A*02:01 presenting the S367-O variant compared to when bound to the S367 cognate peptide by 11.5°C (*p* = 0.01363, **Table S3**). An overlay of the HLA-A*02:01 structures bound to S417 (*24*) and S417-O shows a large difference in the peptide conformation (r.m.s.d. of 1.08 Å, **Fig 2C**). The central part of the peptide shows an inverted conformation of the side chains, with P5-Y and P7-Y of S417-O being buried (**Fig 2D**), and P6-N solvent exposed, while it is the opposite for the S417 peptide (*24*) (**Fig 2C**). The P5-Y and P7-Y side chains in the HLA-A*02:01-S417 structure shows weak electron density, suggesting high mobility (*24*), while it was well defined for the S417-O peptide (**Fig S3C-D**). Surprisingly, the overall stability of the HLA-A*02:01-S417-O complex was significantly lower by 7.4°C (*p* = 0.01392) than that of HLA-A*02:01-S417 (**Table S3**). This might be due to the loss of hydrophobic interactions between P1 residue and W167 from the HLA-A*02:01 (**Fig S4B-C** (*24*)). The P4-D, P6-L and P8-R from the S976 peptide are exposed to the solvent and likely to interact with TCRs (**Fig 2E**). The S976-O epitope is mutated at P6 (L6F), which is solvent exposed, and therefore, the mutation is unlikely to impact on the conformation of the peptide, as per our proposed model (**Fig 2F**), and did not impact on the stability of the pHLA-A*02:01 complexes exhibiting a similar thermal melt point of 50°C within the range of other pHLA-A*02:01 complexes (**Table S3**) (*27*).

**Fig. 2.**
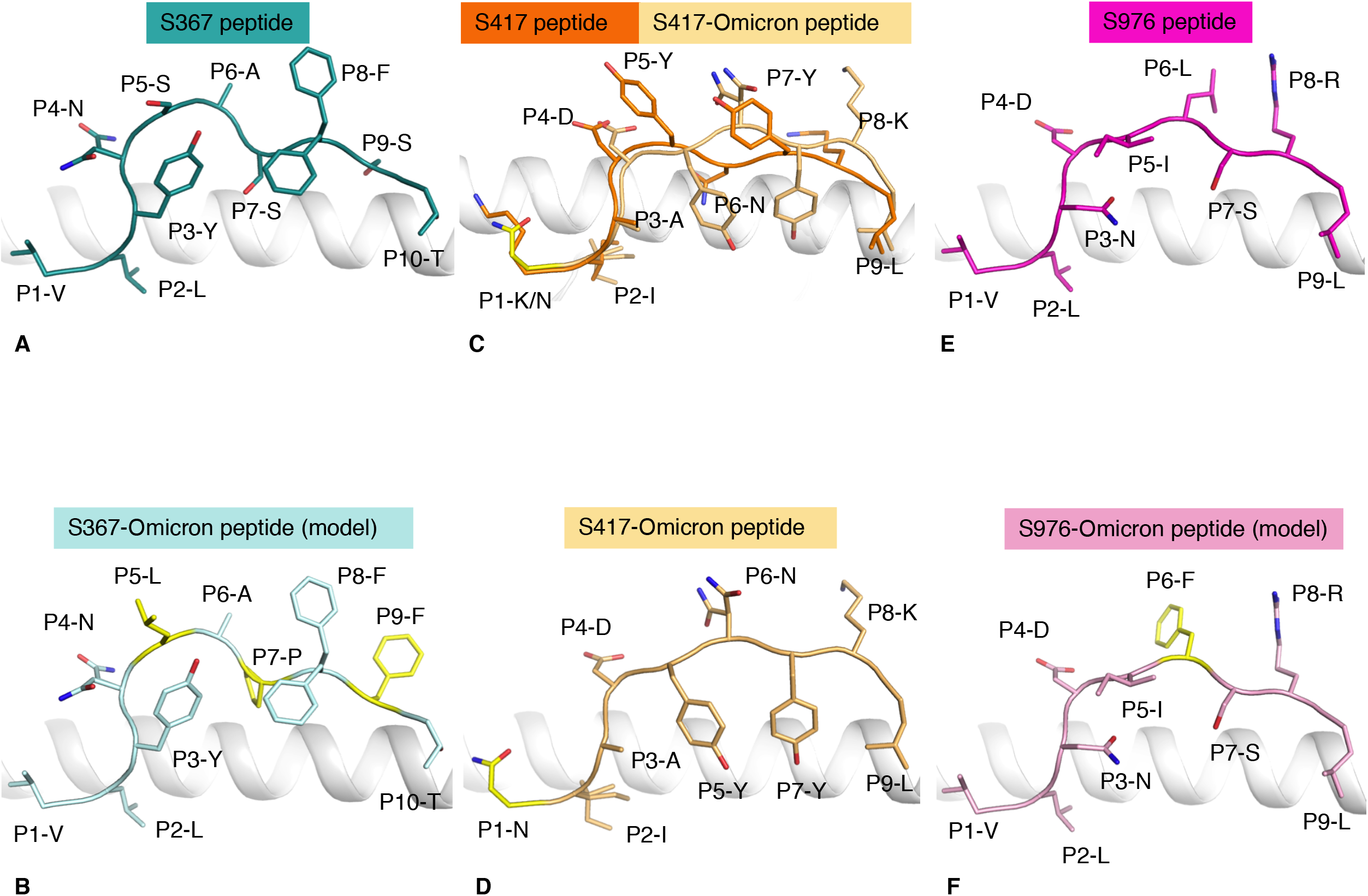
Structural basis for selective T cell cross-reactivity. Crystal structures of the HLA-A*02:01 binding cleft depicted as a white cartoon presenting (**A**) S367 (teal stick), **(B**) S367-O (pale blue stick with Omicron specific mutations in yellow, P5-L, P7-P and P9-F), (**C**) overlay of the crystal structures of the S417 (orange) and S417-O peptides (light orange stick with Omicron specific mutations in yellow, P1-N), (**D**) S417-O (light orange stick with Omicron specific mutations in yellow, P1-N), (**E**) S976 (pink stick), and **(F**) S976-O peptide (light pink stick with Omicron specific mutations in yellow, P6-F).

We next tested whether the Omicron variant peptides, and the cognate peptides, could elicit an *ex vivo* CD8^+^ T cell response in recovered HLA-A*02:01^+^ individuals (**Table S1** and **Fig S5-6**). Responses to S976, S976-O and S367 peptides were only observed in one of three individuals. Two individuals demonstrated a higher CD8^+^ T cell response towards the S417-O and S367-O variants compared to the cognate peptides (**Fig 3A**). The *ex vivo* response towards individual peptides from recovered individuals did not change the median level of CD8^+^ T cell activation over time (**Fig 3A**). We performed an *ex vivo* CD8^+^ T cell responses in PBMC from vaccinated individuals collected after each of the three vaccine doses and 6 months post-1^st^ dose (**Fig 3B-D**). There was a weak but detectable *ex vivo* CD8^+^ T cell response, even after the 1^st^ vaccine dose, towards all peptides, cognate and variants, in at least two of six individuals. The CD8^+^ T cell response did not significantly increase after the 2^nd^ vaccine dose. However, in contrast with nAb levels (**Fig 1B**), T cell responses were similar or stronger 6 months post-1^st^ vaccine dose (**Fig 3B-D**). In addition, T cell responses were increased towards all three Omicron variants compared to the cognate peptides following the booster vaccine (**Fig 3B-D**), with a significant increase observed in the S367-O specific T cell response between the 6 months post-1^st^ dose and the booster vaccine (p < 0.019, **Fig 3D**).

**Fig. 3.**
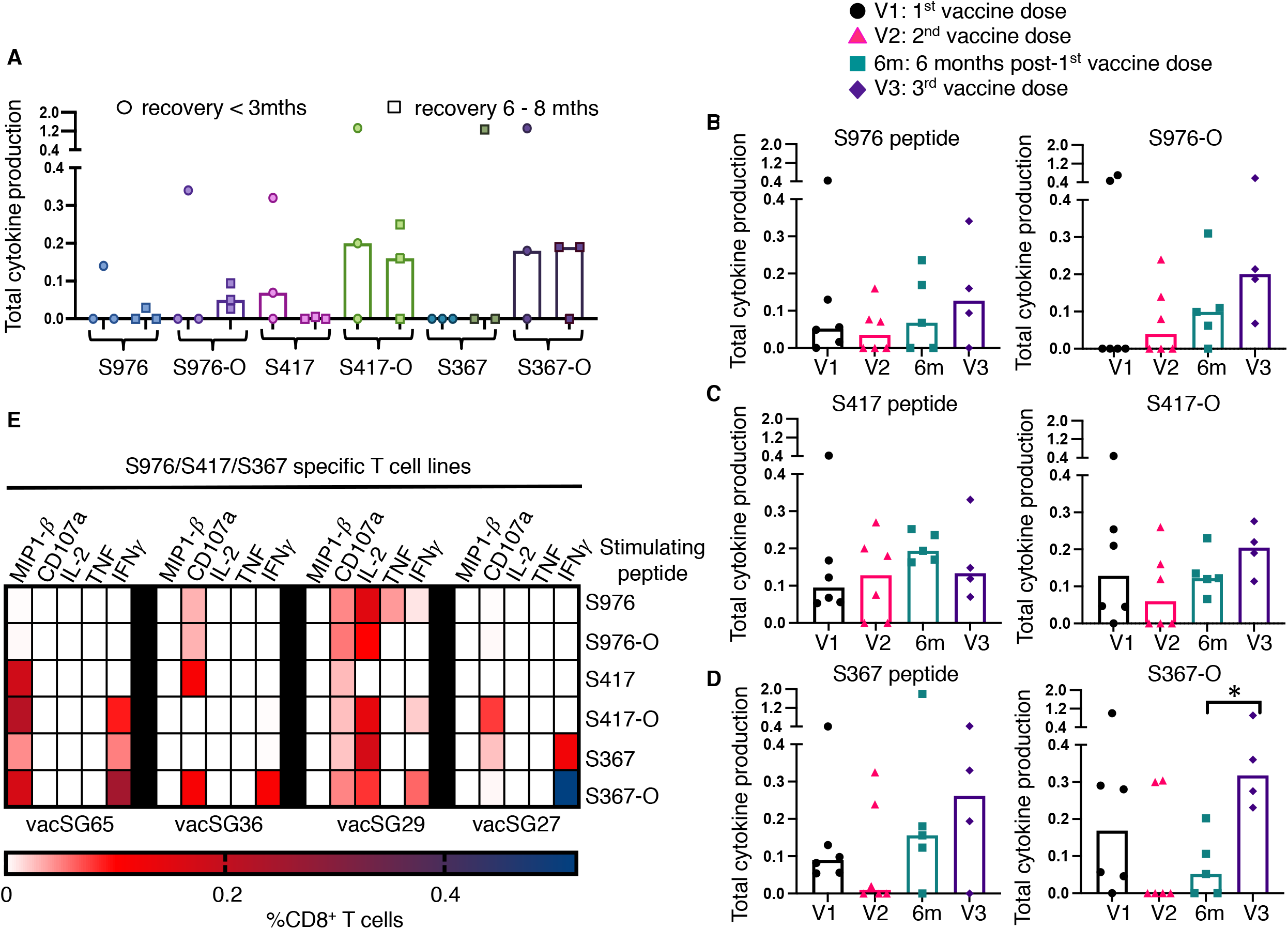
CD8^+^ T cell responses toward spike peptides derived from SARS-CoV-2 and Omicron. PBMCs from HLA-A*02:01^+^ recovered or vaccinated individuals at different recovered or vaccinated time points were stimulated with SARS-CoV-2 and Omicron derived peptides and then assessed for cytokine production **A.** Bar chart depicting the cumulative total of all IFN-γ^+^, all TNF^+^ and all IL-2^+^ CD8^+^ T cells in COVID-19 recovered individuals at <3 months and 3-8 month time points. **B.** Bar chart depicting the cumulative total of all IFN-γ^+^, all TNF^+^ and all IL-2^+^ CD8^+^ T cells towards S976 and S976-O in vaccinated individuals. **C.** Bar chart representing frequency of cytokine response towards S417 and S417-O in vaccinated individuals. **D.** Bar chart representing frequency of cytokine response towards S367 and S367-O in vaccinated individuals. **E.** Heatmap representing the frequency of CD8^+^ IFN-γ, TNF, MIP1β, CD107a and IL-2 producing T cells responding to S367, S417, S976 cognate peptides and S367-O, S417-O, S976-O Omicron variants in vaccinated individuals at 6 months post-1^st^ vaccine dose (n = 4).

To determine the level of T cell activation after 6 months post-1^st^ vaccination *in vitro,* and the T cell cross-reactivity to the Omicron variants, T cell lines were established with the three cognate epitopes together (S367, S417 and S976), and were re-stimulated with single peptide (cognate or Omicron variant). The CD8^+^ T cell polyfunctionality was assessed in samples collected 6 months post-1^st^ dose injection (**Fig 3E** and **Fig S5-S6**). All epitopes were recognized in at least 50% (n=2/4), and up to 100% (n=4/4) of the individuals, and all individuals were able to recognize at least three, and up to all six epitopes, suggesting the presence of memory T cells in all donors. The cytokine profile of the activated CD8^+^ T cells differed between individuals and between epitopes. The S367-O epitope, that carries three mutations, was recognized by all donors and showed a strong cytokine profile with IFNg^+^/CD8^+^ T cell present in all individuals (**Fig 3E**). Interestingly, the level of IFNg-producing CD8^+^ T cells was higher upon presentation of the S367-O peptide than the cognate S367 peptide used to set up the T cell lines in three out of four individuals (**Fig S6**).

Analysis of the Omicron variant epitopes shows that, despite the numerous spike mutations, they can be presented by HLA-A*02:01, although some mutations can impact the peptide conformation. However, the ability of T cells to cross-recognize multiple peptides leads to strong T cell activation against Omicron variants. A stronger T cell activation against the Omicron variants (S417-O and S367-O) compared to the cognate epitopes was observed after 6 months post-1^st^ vaccine dose in some individuals. This could be due to T cell diversity, with some T cells able to engage more effectively with the Omicron variants than cognate epitopes. This diversity could explain the level of T cell activation observed against S417-O and S367-O epitopes even after the 1^st^ vaccine dose alone. This could also reflect that even if the S417 and S417-O peptides are presented in different conformation by HLA-A*02:01, T cells are still able to recognize the Omicron and cognate peptides. It is also possible that some T cells engage with the Omicron variants with higher affinity, or that peptide like S367-O stay on the cell surface longer due to higher stability, leading to higher cytokine production.

In vaccinated individuals, even with only one vaccine dose, CD8^+^ T cells could be activated *ex vivo* against both the cognate and Omicron variant epitopes suggesting provision of early protection against the SARS-CoV-2 Omicron. In addition, the 2^nd^ vaccine dose increased the nAb levels, spike-specific memory B and T cells detected in peripheral blood. The nAb response slowly decreased over time, and after 6 months was comparable to the response levels detected after the 1^st^ vaccine dose, still providing a level of protection, albeit less. Interestingly, recall of spike-specific T cell responses in PBMCs were increased following the booster vaccine, higher than the levels detected following the 2^nd^ vaccine dose.

The presence of CD8^+^ T cells able to produce cytokines towards the Omicron variants, in the context of HLA-A*02:01, in combination with cross-reactive nAb (*4–8*) and long lived memory B cells show that the current vaccines are likely to provide a level of protection against Omicron. In addition, booster vaccination not only increases the nAb levels to Omicron (*8*), it induces a stronger T cell response, especially against the triple mutated S367-Omicron epitope. This evidence of memory T cell responses towards the Omicron variants in individuals previously infected with non-Omicron variants, or that have received vaccines based on ancestral strain of SARS-CoV-2, could be critical in protecting people from severe COVID-19 if infected with this highly transmissible variant.

## Acknowledgments

The authors would like to thank Queensland Health Forensic & Scientific Services, Queensland Department of Health who provided the SARS-CoV-2 isolate QLD02; La Trobe University Facilities (Imaging platform with Flow cytometers, Crystallisation Facility); VTIS and CareDx for HLA Typing; MX team for assistance at the Australian Synchrotron. This project obtained samples for the COVID-19 Biobank based at the Alfred Hospital and acknowledge all its investigators and the participants who contributed samples. We acknowledge the research coordinators in the clinical research unit at the Alfred Hospital. The authors would also like to thank all the participants who took part in our study.

## Author contributions

Conceptualization: A.T.N., C.S., D.S.M.C., S.G.; Methodology: A.T.N., C.S., D.S.M.C., Z.W.M.T., M.D.O., L.C., K.Y.C., B.G., D.J., E.J.G., C.Smith., J.M., K.L.G-J., P.J.B., K.R.S., S.G.; Investigation: A.T.N., C.S., D.S.M.C., Z.W.M.T., K.Y.C., M.D.O., L.C., L.D.M., A.R.T., R.W., D.J., Y.M.A., S.G.; Visualization: A.T.N., C.S., D.S.M.C., Z.W.M.T., L.C., C.A.L., S.G.; Reagents: K.E.L., B.G., C.Smith., J.M.; Funding acquisition: C.Smith., J.M., K.L.G-J., P.J.B., K.R.S., S.G.; Project administration: S.G.; Supervision: C.Smith., J.M., K.L.G-J., P.J.B., K.R.S., S.G.; Writing – original draft: S.G.; Writing – review & editing: all authors.

## Data and materials availability

All data are available in the main text or the supplementary materials.

