## Supplementary figures and images for "COVID-19 vaccine booster induces a strong CD8^+^ T cell response against Omicron variant epitopes in HLA-A*02:01^+^ individuals"

### Supplemental figures

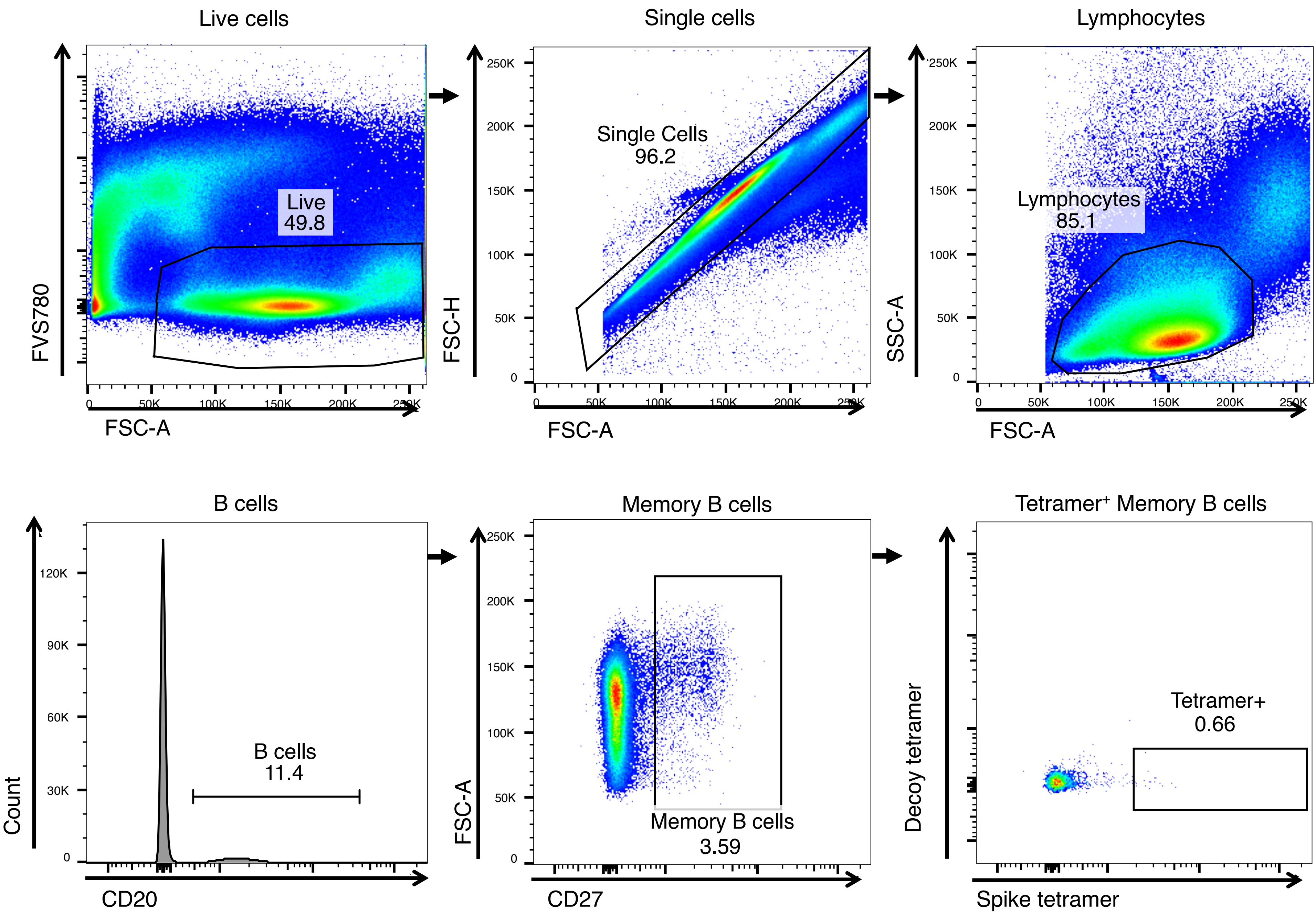

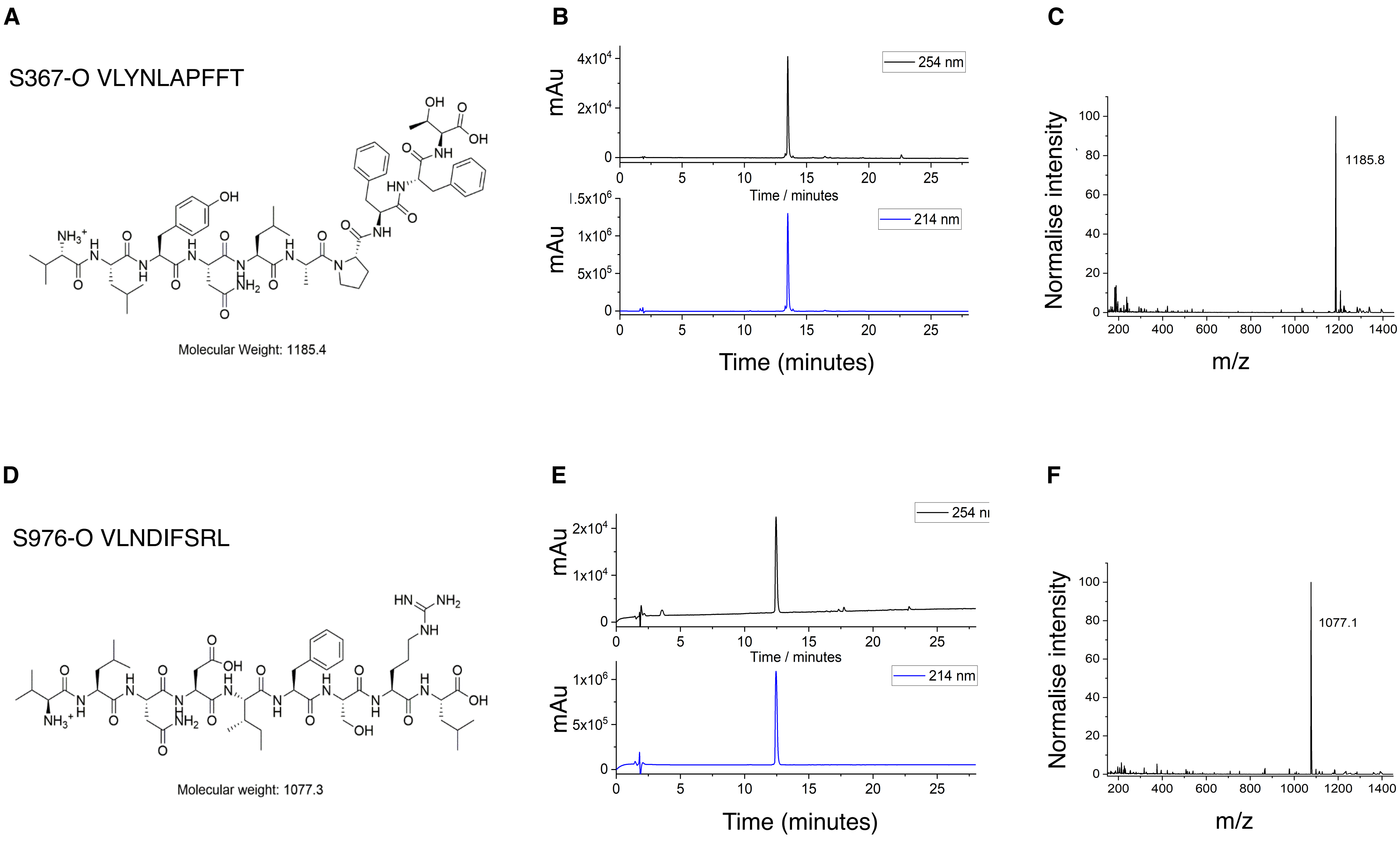

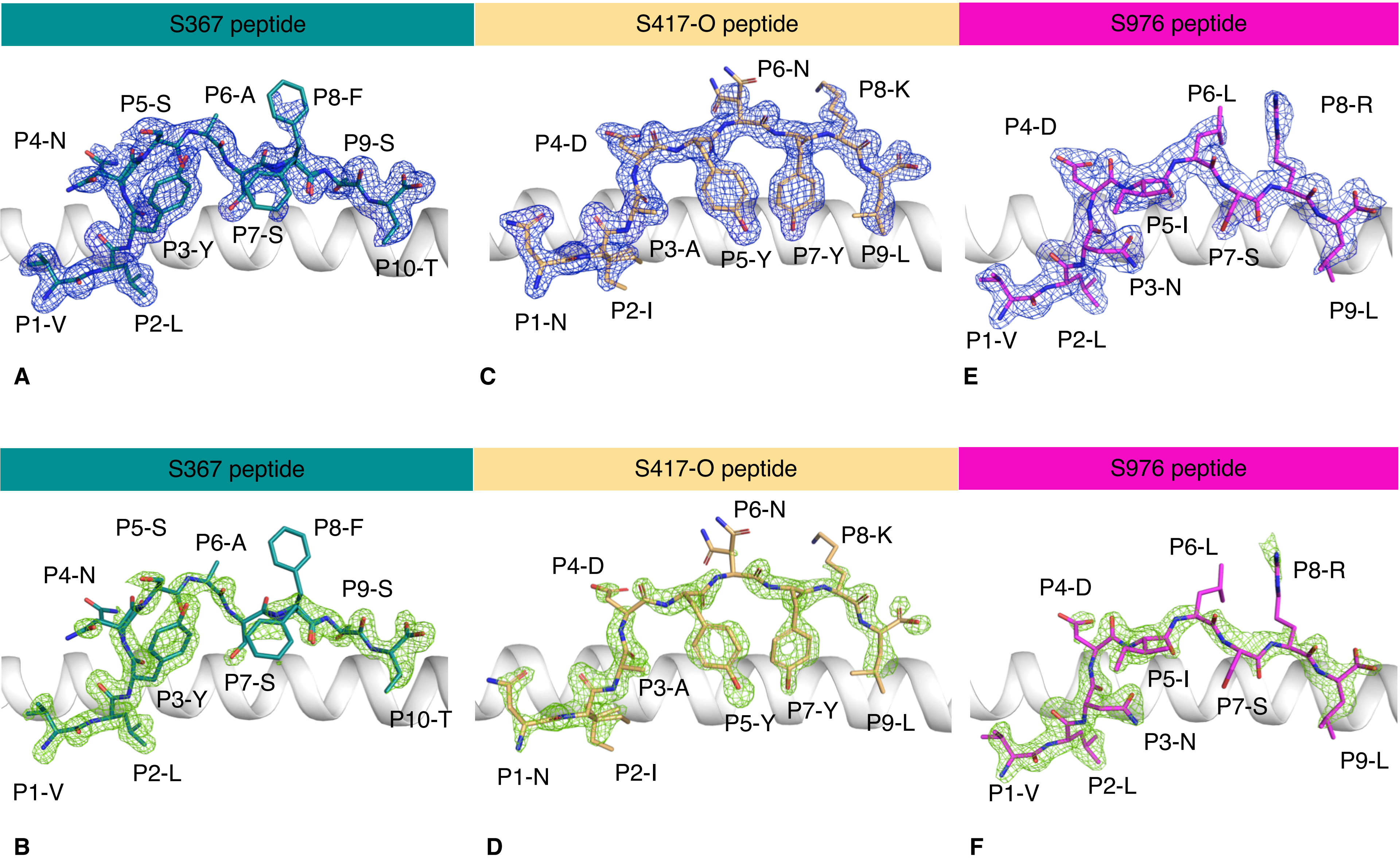

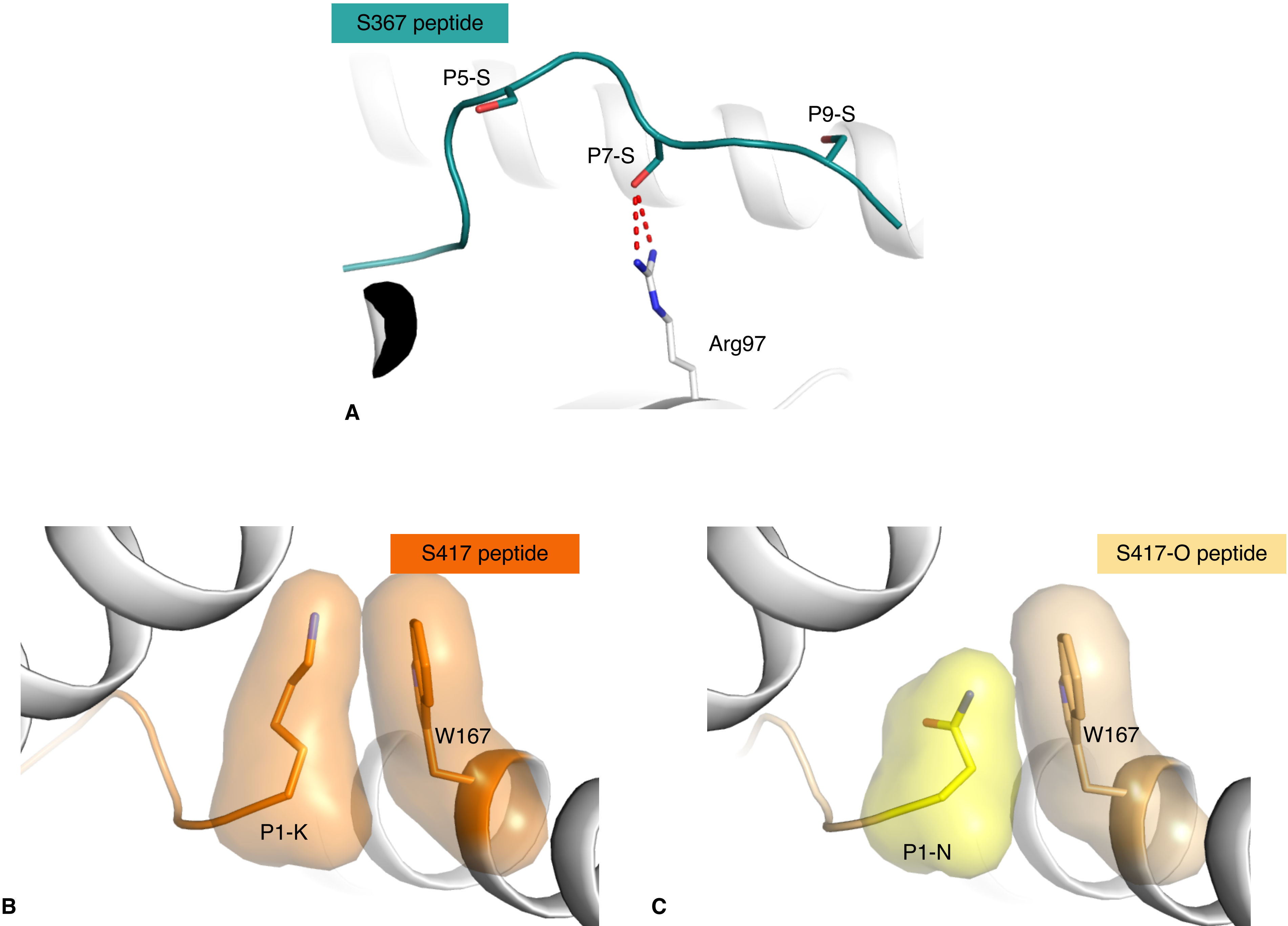

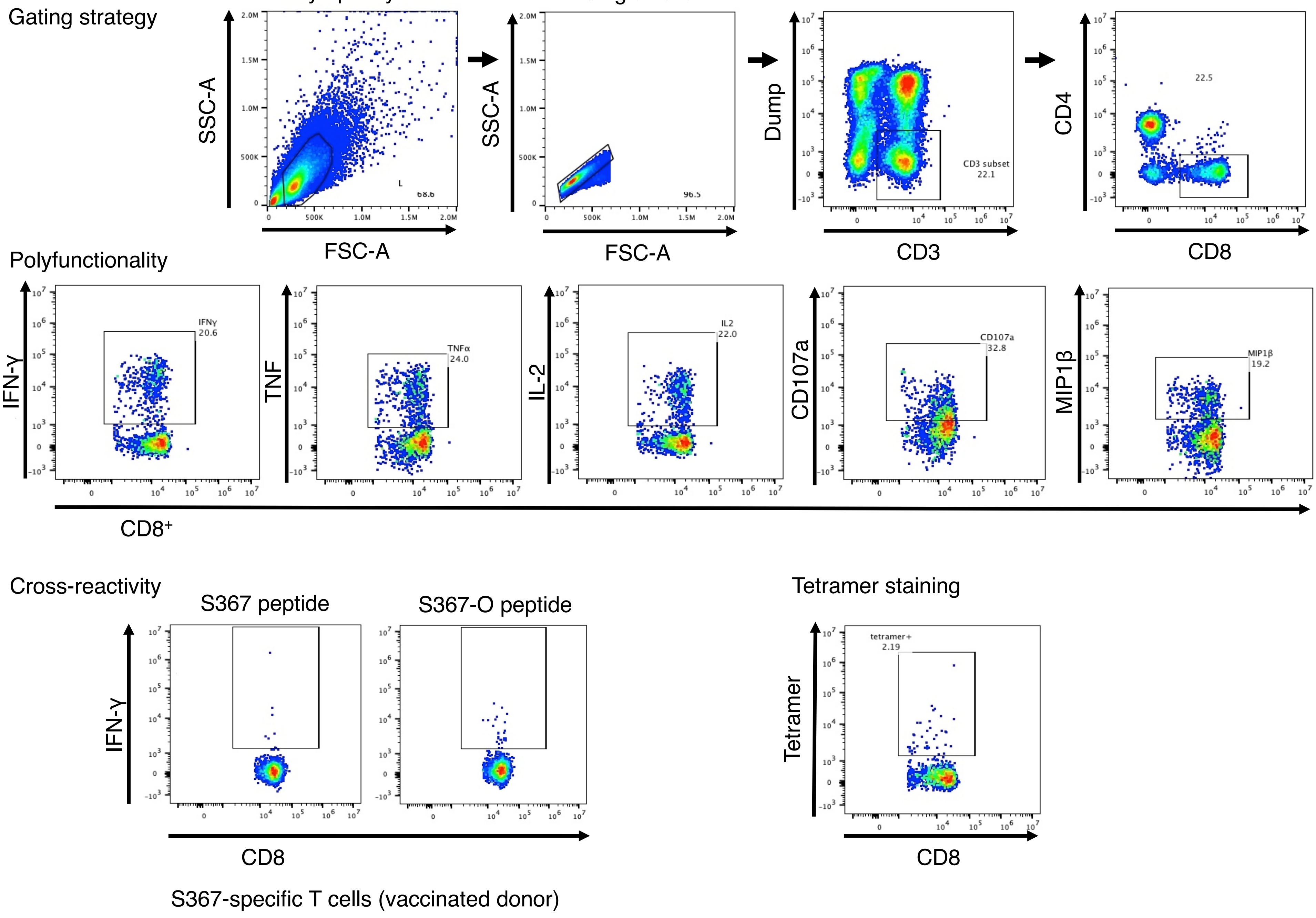

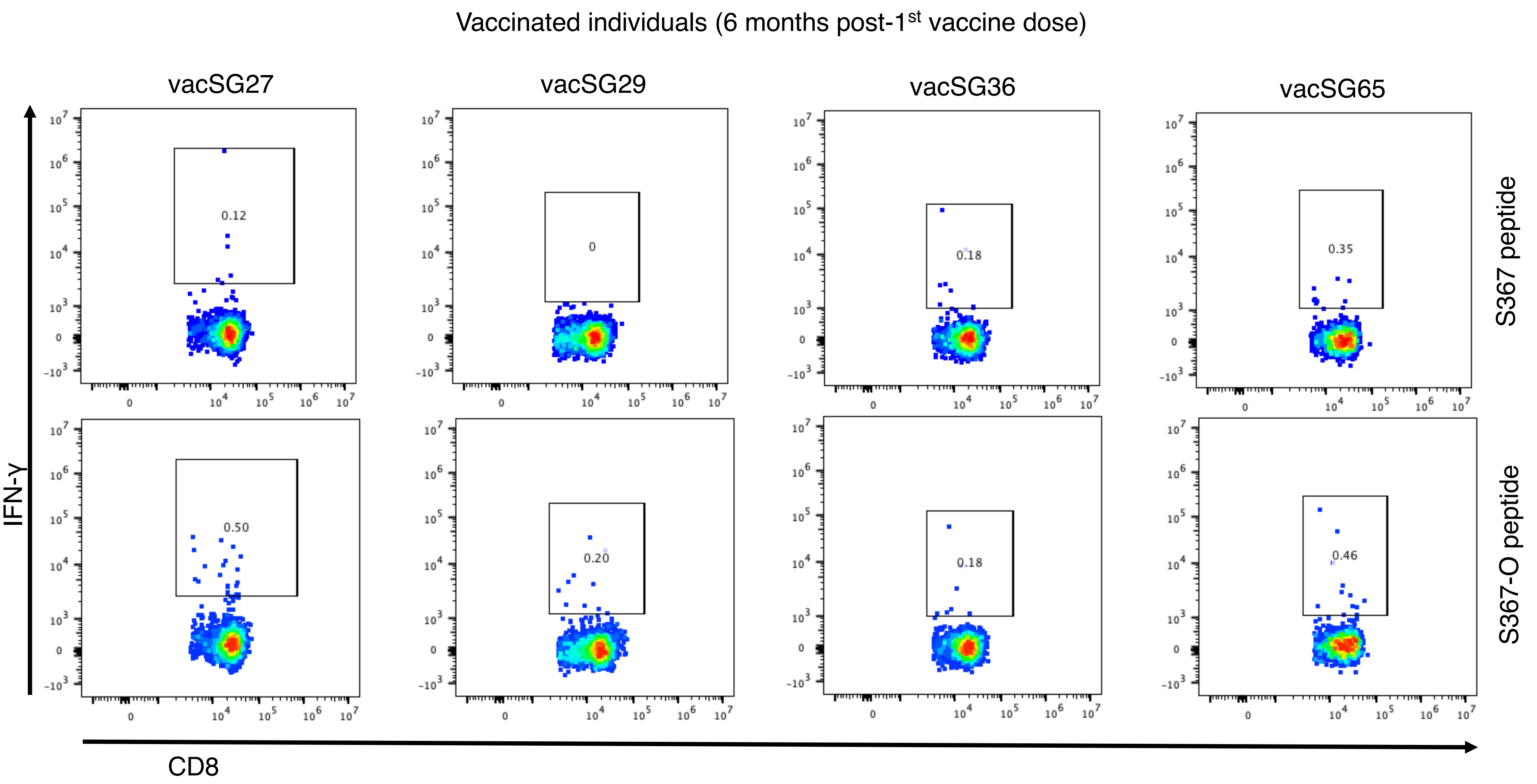
