## Supplemental data for "COVID-19 vaccine booster induces a strong CD8^+^ T cell response against Omicron variant epitopes in HLA-A*02:01^+^ individuals"

### Supplementary Materials

### Materials and Methods

#### COVID-19 recovered participant samples

Individuals with a PCR confirmed diagnosis of COVID-19 were enrolled to donate additional samples of blood and detailed clinical information (Alfred Health Human Research and Ethics Committee Number 182/20).

#### COVID-19 vaccinated participant samples

Volunteer participants were given a consent form, explanatory statement of the research and a questionnaire (La Trobe University Human Research and Ethics Committee Number HEC21097). PBMCs were separated from whole blood using density gradient centrifugation and used fresh or cryogenically stored until use.

#### Neutralising Antibody assay

Neutralising antibodies to SARS-CoV-2 (hCoV-19/Australia/QLD02/2020; QLD/02) were assessed as described previously *(14)*. SARS-CoV-2 was kindly provided by Queensland Health Forensic and Scientific Services. All work with infectious SARS-CoV-2 was performed under BSL3 conditions. Briefly, Vero cells were cultured in 96 well plates. Convalescent serum harvested from COVID-19-recovered patients was heat-treated at 56°C for 1 hour. The diluted sera was then serial diluted with minimum essential medium (MEM) (GIBCO) supplemented with 2% FCS. In physical containment 3 settings, the diluted sera was incubated with SARS-CoV-2 (QLD/02; MOI 0.1) for 1 hour at room temperature. The serum-virus mixture was transferred to the cultured Vero cells and further incubated for 1 hour at room temperature for infection. The inoculum was then removed and replaced with MEM (GIBCO) supplemented with 2% FCS (GIBCO). The cells were incubated at 37°C for 48 hours. The cells were fixed with 10% formaldehyde for 24 hours. Cells were permeabilised with PBS containing 0.1% Trition-X-100 (Sigma) for 15mins at room temperature. The plates were blocked using PBS (Sigma) supplemented with 0.1% Tween-20 (Sigma) and 3% skim milk for 1 hour at room temperature. The cells were primarily stained with rabbit anti-NP (1:5000)(Sino Biological) for 1 hour at room temperature. The plates were washed with PBS (Sigma) supplemented with 0.1% Tween-20 (Sigma). The cells were stained using goat anti-rabbit immunoglobulin horse-radish protein (HRP) (Thermo Fischer) (1:5000) for 1 hour at room temperature. The plates were washed with PBS (Sigma) supplemented with 0.1% Tween-20 (Sigma). The plates were incubated with 2,2′-azino-bis(3-ethylbenzothiazoline-6-sulfonic acid (ABTS) (Thermo Fischer) substrate for color development and incubated for 20 minutes. Measurements were done at 405 nm.

#### Peptide Synthesis and Purification

Peptides VLYNLAPFFT (S367-O) and VLNDIFSRL (S976-O) were synthesised using a standard fluoren-9-ylmethoxycarbonyl (Fmoc) automated solid-phase peptide synthesis method with Wang resin (100−200 mesh, 1.24 mmol/g) utilising microwave (MW)-assisted heating on a Biotage® Initiator+ Alstra peptide synthesiser (**Fig S6**). Prior to use the Wang resin was swollen in DMF for 2 hours. The Fmoc-amino acids and HCTU/HOBt/DIPEA (4 eq.) dissolved in DMF were then added sequentially to the resin and the reactions were carried out with MW heating at 60 °C for 20 min. The Fmoc protecting groups were removed using piperidine (20% in DMF) at RT for 20 minutes. Peptides were cleaved from the resin using a solution of 95% trifluoroacetic acid (TFA), 5% triisopropylsilane (TIPS) for 3 h. The acid was evaporated, and the crude peptide was purified using RP-HPLC using a Shimadzu HPLC fitted with two Shimadzu LC-20AD pumps, a SIL-20AHT autosampler, a SPD-M20A photodiode array detector and a FRC-10A fraction collector and an Onyx Monolithic C18, 100 × 10 mm semi-preparative column with a gradient of 0.1% TFA in water (solvent A) and acetonitrile (solvent B) over 30 min. Purified peptides were lyophilized and stored at −20 °C. Purity was confirmed at >95% in each case by analytical HPLC and structures were confirmed using electrospray ionization mass spectrometry (**Fig S6**).

**Generation of peptide-specific CD8$^+$ T cell lines**

CD8$^+$ T cell lines were generated as previously described (*29*). Briefly, PBMCs from COVID-19 recovered or vaccinated individuals were thawed and split 1:3. 1/3 of the PBMCs were pulsed with 2μM of SARS-CoV-2 overlapping peptide pools or 10μM individual peptides and incubated for 90 minutes at 37°C. The stimulated cells were then washed twice with RF10 media (RPMI-1640 [Thermofisher, Scoresby, Australia] supplemented with 2 mM MEM nonessential amino acid solution [Thermofisher], 100 mM HEPES [Thermofisher], 2 mM L-glutamine [Thermofisher], penicillin/streptomycin [Thermofisher], 50 mM 2-ME [Sigma] and 10% heat-inactivated fetal calf serum [FCS; Scientifix]); and added together with the 2/3 of unpulsed PBMCs. T cell lines were cultured for 10 days in RF10 media. The cultures were supplemented with 20 IU IL-2 (BD Biosciences, Melbourne, Australia) 2–3 times weekly. CD8$^+$ T cell lines were freshly harvested and used for subsequent assays.

**Intracellular cytokine staining (ICS) assay**

The *ex vivo* T cell assay was performed with $10^6$ PBMCs from HLA-A*02:01$^+$ individuals were thawed, stimulated with 10μM of individual peptides (Genscript, Hong Kong, China) and incubated for 10 hours in RF10 media (described above) supplemented with 20 IU IL-2 (Peprotech), in the presence of GolgiPlug, GolgiStop and anti-CD107a-AF488-FITC (BD Biosciences).

The peptide-specific CD8$^+$ T cell lines were used for *in vitro* intracellular cytokine staining assay and was performed as previously described *(14)*. In summary, CD8$^+$ T cell lines were stimulated with cognate peptide pools or 1μM of individual peptides (Genscript, Hong Kong, China) and incubated for 5 hours in the presence of GolgiPlug, GolgiStop and anti-CD107a-AF488 (all BD Biosciences).

In both cases, following stimulation, cells were surface stained for 30 minutes with anti-CD8-PerCP-Cy5.5 (BD Biosciences/eBioscience), anti-CD4-BUV395 (BD Biosciences), anti-CD14-APCH7, CD19-APCH7 and Live/Dead Fixable Near-IR Dead Cell Stain (Life Technologies, Melbourne, Australia). Following incubation, cells were fixed and permeabilised for 20 minutes using BD Cytofix/Cytoperm solution (BD Biosciences) then intra-cellularly stained with anti-IFN-γ-BV421, anti-TNF-PE-Cy7, anti-MIP1β-APC and anti-IL-2-PE (BD Biosciences) for a further 30 minutes. Cells were acquired on a CytoFLEX (Beckman Coulter). Post-acquisition analysis was performed using FlowJo software (version 10.7.1, BD Biosciences). Cytokine detection levels identified in the no-peptide control condition were subtracted from the corresponding test conditions in all summary graphs to account for non-specific, spontaneous cytokine production. Due to the low frequency of cytokine production by CD8$^+$ T cells *ex vivo*, and variability in cytokine preference between donors, data is shown as a cumulative total of all IFN-$\gamma^+$, all TNF$^+$ and all IL-2$^+$ CD8$^+$ T cells.

**Spike protein expression and purification**

The construct of the Spike protein used to generate tetramer contain the Hexapro sequence *(30)*, which is a mutated version of the spike to increase protein production in mammalian cells as well as stabilizing the pre-fusion conformation of the spike. We then added a BirA tag and a 6 His tag to enable biotinylation (required for tetramerization) and purification, respectively. The final construct was codon optimized and synthesized (Genscript, Hobg-Kong, China) and sub-cloned into a pHLsec vector to be expressed in HEK293T cells. The plasmid was used to transform XL-1 Blue chemically competent cells, from which overnight culture were grown in LB broth at 37°C for 14 hours. High quality DNA was extracted from the cells using a plasmid gigaprep kit (Cat: D4204, Zymo Research, Victoria, Australia). HEK293T cells were grown in a $CO_2$ incubator in DMEM *(Dulbecco's Modified Eagle Medium)* media containing 10% Fetal bovine serum (Cat: SFBS, Bovogen Biologicals, New Zealand) and 1% Penicillin-Streptomycin-Glutamine (Cat: 10378016, Thermofisher, Victoria, Australia). The cells were grown for 3 days in 5-layer Multilayer flasks (Cat: 353144, Corning, Victoria, Australia) until 70% confluency. Purified DNA was incubated for 20 minutes with Polyethylenimine (PEI) (Cat: 408727-100ML, Sigma, Victoria, Australia) at a ratio of 1:3. During the incubation, the 10% FBS supplemented media was removed from the flasks and replaced with DMEM containing 1% FBS, 5% cocktail of supplements referred to as Mastermix (DMEM with 1% Glutamax, 1% sodium pyruvate, 1% hepes, 1% NEAA, 0.000004% betamercaptoethanol). Following the incubation period, the DNA:PEI mix was added to the cell culture flasks, and placed back in the $CO_2$ incubator for 7 days. Following 7 days of protein expression, the supernatant was harvested and precipitated with 1mM Nickel Chloride and 5mM Calcium chloride for 20 minutes, then centrifuged for 10 min at 6000g. It was then further filtered with glass microfibre filters (Cat: FF0483-047, MC Scientific, Victoria, Australia) and cellulose acetate membrane filters (Cat: 11104--47------N, Sartorius, Victoria, Australia) using a Solvac filter holder (Cat: 4020, Pall, Victoria, Australia). It was then adjusted to contain 20mM Imidazole pH 8 and bound onto a Histrap HP column (Cat: GE17-5248-02, Cytiva, Victoria, Australia) using a peristaltic pump (Cat: 56300551, Cytiva, Victoria, Australia), and eluted with a gradient of Imidazole (stock at 1M pH 8). The spike protein was eluted with ~300mM Imidazole pH 8, concentrated and buffer exchange to be applied onto a HiLoad 16/600 Superdex size exclusion column (Cat: 28989335, Cytiva, Victoria, Australia). The protein was run on a SDS-page gel to confirm purity and then biotinylated and stored at -20°C.

**Spike protein tetramer conjugation**

The Decoy tetramer was prepared by conjugating the core fluorochrome SA-PE to DL650 (Sigma-Aldrich) according to the manufacturer's protocol for 60 minutes at room temperature. The free DL650 was removed by centrifugation in a 100-kD molecular weight cut off Amicon Ultra filter (Millipore). The SA-PE*DL650 complex concentration was calculated by measuring the absorbance of PE at 566nm using a NanoDrop ND-1000 spectrophotometer (Thermo Fisher Scientific). The SA-PE*DL650 complex was then incubated with 10-fold molar excess of biotinylated HLA-A*02:01-M1$_{58}$ for 30 minutes at room temperature. The solution was diluted to 1μM based on the absorbance of PE at 566 nm (divided by the extinction coefficient = 1.96 $cm^{-1}\mu M^{-1}$). The COVID-19-specific B cell tetramer was prepared by conjugating the biotinylated

HexaPro spike protein to SA-PE (Prozyme PJRS25) at a concentration ratio of 4:1 respectively. Following addition of SA-PE to the biotinylated HexaPro spike protein, the mixture was incubated in the dark at room temperature for 3 hours (with gentle mixing every 30 minutes). Following this, the mixture was stored overnight at 4°C in the dark. The tetramer fraction was then centrifuged in a 100-kD molecular weight cutoff Amicon Ultra filter (Merck Millipore). Because the ratio of SA/fluorochrome is roughly 1:1, the concentration of tetramer was calculated by measuring the absorbance of PE at 566 nM (divided by the extinction coefficient = 1.96 $cm^{-1}\mu M^{-1}$). The tetramer was diluted to a concentration of 1μM and stored at 4°C in the dark. During flow cytometric analysis, cells that bind to the HexaPro spike protein tetramer as well as the Decoy SA-PE*DL650 tetramer were considered Decoy-specific and excluded from further analysis.

**Antibody staining and flow cytometry for the B cells**

Staining for flow cytometry analysis was performed using cryo-preserved PBMCs. PBMCs were stained with Fixable Viability Stain 780 (FVS780) (1:1000, BD Biosciences) diluted in PBS 2% FCS for 15 minutes at room temperature. Cells were washed with PBS 2% FCS and then stained with CD20-AF700 (2H7, 1:200), CD27-V450 (M-T271, 1:50), CD38-FITC (90, 1:25), IgD-BV510 (IA6-2, 1:50), CD10-PE-Cy7 (HI10a, 1:50), HexaPro spike protein Tetramer-PE (1:50) and HLA-A2-$M1_{58}$ Decoy Tetramer-PE*DL650 (1:100) (all BD Biosciences except for the tetramers which were made in house) diluted in PBS 2% FCS for 20 minutes on ice. Cells were washed with PBS 2% FCS and fixed for 1 hr using Cytofix fixation buffer (BD Biosciences), then washed with PBS 2% FCS and resuspended in PBS 2% FCS. Cells were analysed using a LSRFortessa X-20 (BD Biosciences) and flow cytometry data was analysed using FlowJo™ v10.8 software (BD Biosciences).

**Protein expression, refold and purification of pHLA complexes**

DNA plasmids encoding HLA-A*02:01 α-chain and β-2-microglobulin were transformed separately into a BL21 strain of *E. coli*. Recombinant proteins were expressed individually, where inclusion bodies were extracted and purified from the transformed *E. coli* cells as per previously described in details (*31*). Soluble pHLA-A*02:01 complexes were produced by refolding 30 mg of HLA-A*02:01 α-chain with 10 mg of β-2-microglobulin and 5 mg of peptide (Genscript) into a buffer of 3M Urea, 0.5 M L-Arginine, 0.1 M Tris-HCl pH 8.0, 2.5 mM EDTA pH 8.0, 5 mM glutathione (reduced), 1.25 mM glutathione (oxidised). The refold mixture was dialysed into 10 mM Tris-HCl pH 8.0 and soluble pHLA complexes were purified using anion exchange chromatography using a HiTrapQ column (GE Healthcare).

**Differential scanning fluorimetry**

Thermal stability assay was performed by Differential Scanning Fluorimetry carried out in ViiA 7 real-time PCR system (Thermofisher), where pHLA samples were heated from 25 to 95˚C at a rate of 1˚C/min in 0.5˚C steps. The excitation and emission channels were set to the TAMRA reporter (x3m3 filter) with excitation of ~550 nm and detection at ~587 nm. The experiment was performed at two concentrations of pHLA (5 μM and 10 μM) in duplicate. Each sample was dialysed in 10mM Tris-HCl pH 8.0, 150mM NaCl and contained a final concentration of 10X SYPRO Orange Dye. Fluorescence intensity data was normalised and plotted using GraphPad Prism 9 (version 9.0). The Tm value for a pHLA is equal to its the temperature when 50% of maximum fluorescence intensity is reached, which approximately equals to 50% of unfolded protein and summarized in **Table S3**.

**Crystallization and structural determination**

Crystallisation of pHLA complexes were grown via sitting-drop, vapour diffusion at 20˚C. The protein:reservoir drop ratio is 1:1, at a concentration of 3 mg/mL in 10 mM Tris-HCl pH 8, 150 mM NaCl. Crystals of HLA-A*02:01 in complex with S367, S417-O and S976 were grown in 20% P3350; 0.2 M Li Acetate; 14% P3350; 0.1M NaFlu, 1mM $CdCl_2$, and 13% P3350; 0.1M NaFlu; 2% EG; 1 mM $CdCl_2$, respectively. Protein crystals were soaked in a cryoprotectant solution containing mother liquor solution with the PEG3350 concentration increased to 30% (w/v) and then flash-frozen in liquid nitrogen. The data were collected on the MX2 beamline at the Australian Synchrotron, part of ANSTO, Australia (*32*). The data were processed using XDS (*33*) and the structures were determined by molecular replacement using the PHASER program (*34*) from the CCP4 suite (*35*) with a model of HLA-A*02:01 without the peptide (derived from PDB ID: 7KGS (*27*)). Manual model building was conducted using COOT (*36*) followed by refinement with BUSTER (*37*). The final models have been validated and deposited using the wwPDB OneDep System and the final refinement statistics, PDB codes are summarized in **Table S2**. All molecular graphics representations were created using PyMOL.

**Quantification and statistical analysis**

GraphPad Prism 9.3.0 (San Diego, CA) was used to generate the graphs from the analysis and to perform statistical analysis. Statistical comparison between column of data was performed using the Wilcoxon test, and $p < 0.05$ was considered statistically significant.

### Supplementary Figure

**Figure S1. Gating strategy for spike-specific memory B cells**

PBMCs were isolated from COVID-19 recovered and vaccinated donors. Representative gating for the identification of B cells (top panel). Representative gating for the identification of spike protein-specific memory B cells $CD20^{+}CD27^{+}Tetramer^{+}Decoy^{neg}$ memory B cells (bottom panel). The decoy tetramer was HLA-A*02:01-$M1_{58}$ tetramer.

**Figure S2. HPLC traces for the S367-O and S976-O peptide synthesis**

**A**. Formula of the S367-O peptide with its molecular weight. **B**. Analytical HPLC chromatogram (254 nm and 214 nm) obtained for a purified sample of S367-O (VLYNLAPFFT). The peptide elutes at 13.49 minutes. **C**. Positive-ion ESI-mass spectrum for the S367-O peptide (VLYNLAPFFT). $[M+H]^{+}$=1085.4. **D**. Formula of the S976-O peptide with its molecular weight. **E**. Analytical HPLC chromatogram (254 nm and 214 nm) obtained for a purified sample of S967-O (VLNDIFSRL). The peptide elutes at 12.45 minutes. **E**. Positive-ion ESI-mass spectrum for the S967-O peptide (VLNDIFSRL). $[M+H]^{+}$=1077.3.

**Figure S3. Electron density maps for the crystal structures of SARS-CoV-2 and Omicron variant Spike peptides in complex with HLA-A*02:01**

Density map for the structures of HLA-A*02:01 (white cartoon) binding to Spike-derived SARS-CoV-2 or Omicron variant peptides. (**A-B**) SARS-CoV-2 S367 (teal), (**C-D**) S417-O (light orange) and (**E-F**) S976 (magenta). The peptides are represented as stick, with the electron density map after refinement shown by a blue 2Fo-Fc map contoured at 1 sigma on the top panels (**A, C, E**). The Fo-Fc omit map (after molecular replacement without peptide) is contoured at 3 sigma and coloured in green on the bottom panels (**B, D, F**).

**Figure S4. Peptide and HLA-A*02:01 interaction**

The HLA-A*02:01 is represented as white cartoon on all panels. (**A**) Interaction of the S367 peptide (teal) with the HLA-A*02:01 molecule Arg97 (white stick) by a hydrogen bond (red dashed line) with the P7-S residue. (**B-C**) Interaction of the P1 residue of the S417 peptide (**B**) or S417-O peptide (**C**) with the W167 residue from the α2-helix of the HLA-A*02:01 molecule. The P1-K, P1-N, and W167 are presented as stick and surface to show the interaction between the peptide and HLA residues. As per panel B the P1-N mutated in S417-O (**C**) is coloured in yellow.

**Figure S5. Gating strategy for T cell assay**

Peptide-specific $CD8^{+}$ T cells were derived by stimulating PBMCs from COVID-19 recovered and vaccinated donors with various peptides as indicated throughout. $CD8^{+}$ T cell responses were assessed by either tetramer staining or function in an ICS assay. Representative gating for the identification of $CD8^{+}$ T cells in all assays (top panel). Representative gating for the assessment of polyfunctionality (middle panel) or cross-reactivity (bottom panel, left).

Representative flow cytometry plot of the identification of tetramer$^+$ populations (bottom panel, right).

**Figure S6. T cell activation in vaccinated individuals upon S367 and S367-O peptides presentation**

S367-specific CD8$^+$ T cells were expanded *in vitro* by stimulating PBMCs from HLA-A*02:01 vaccinated donors (n=4) and and cultured for 10 days with IL-2. Specificity and cross-reactivity was assessed by stimulating CD8$^+$ T cell lines with either S367 or S367-O peptides in an ICS assay. Representative FACS plots of IFN$\gamma^+$ production in the ICS assay (n=4) is shown.

**Supplementary Tables S1 to S3**

**Table S1. Donor Information.**

| **ID Recovered** | **Post recovery / COVID Severity** | **sex** | **age** | **HLA-A** | **HLA-B** |
|---|---|---|---|---|---|
| Q015 | 1.5 month / nil | F | 30 | 02:01 | 35:03 |
| ALF0011 | 2 months / NA | F | 42 | 02:01, 11:01 | 27:05, 38:01 |
| ALF0129 | 2 months / NA | M | 38 | 02:01, 26:01 | 15:01, 38:01 |
| ALF0209 | 3 months / NA | F | 62 | 02:01, 24:03 | 08:01, 38:01 |
| ALF0167 | 3 months / NA | F | 26 | 02:01, 03:01 | 14:02, 40:01 |
| ALF0149 | 3 months / NA | M | 52 | 02:01, 03:01 | 44:02, 55:01 |
| Q042 | 3 months / mild | M | 75 | 02:01, 03:01 | 13:02, 40:01 |
| ALF0170 | 3 months / NA | F | 30 | 02:01 | 44:02, 51:01 |
| Q062 | 4 months / mild | F | 22 | 02:01 | 08:01, 18:01 |
| coSG21 | 4 months / mild | F | 51 | 02:01 | 08:01, 37:01 |
| ALF0024 | 6 months / mild | F | 23 | 02:01, 11:01 | 07:02, 35:01 |
| ALF0101 | 7 months / mild | M | 26 | 02:01 | NA |
| coSG5 | 8 months / mild | F | 55 | 02:01, 24:02 | 35:01, 40:01 |
| **ID Vaccinated** | **Vaccine used** | **sex** | **age** | **HLA-A** | **HLA-A** |
| vacSG31 | Pfizer | M | 29 | 02:01, 31:01 | 18:01, 35:01 |
| vacSG35 | AstraZeneca | M | 60 | 01:01, 02:01 | 08:01, 44:02 |
| vacSG7 | AstraZeneca | F | 56 | 01:01, 02:01 | 08:01, 44:03 |
| vacSG40 | AstraZeneca | F | 66 | 02:01, 24:02 | 08:01, 44:02 |
| vacSG21 | Pfizer | M | 23 | 01:01, 02:01 | 08:01, 49:01 |
| vacSG29 | Pfizer | M | 25 | 02:01, 03:01 | 07:02, 35:01 |
| vacSG45 | Pfizer | M | 33 | 02:01, 24:02 | 07:02, 14:02 |
| vacSG51 | Pfizer | M | 26 | 02:01, 03:02 | 38:01, 50:01 |
| vacSG62 | Pfizer | F | 23 | 02:01, 24:02 | 15:01, 48:01 |
| vacSG20 | Pfizer | M | 70 | 02:01, 11:01 | 41:02, 52:01 |
| vacSG27 | Pfizer | F | 76 | 02:01, 03:01 | 07:02,14:02 |
| vacSG36 | Pfizer | F | 61 | 02:01, 11:01 | 35:01, 44:02 |
| vacSG43 | AstraZeneca | F | 74 | 01:01, 02:01 | 35:01, 44:02 |

NA: not available; M: male; F: female; age in years; AstraZeneca: ChAdOx1 nCoV-19 vaccine; Pfizer: BNT162b2 Pfizer-BioNTech COVID-19 vaccine.

**Table S2. Data Collection and Refinement Statistics**

| **Data Collection Statistics** | **HLA-A*02:01-S417-O** | **HLA-A*02:01-S367** | **HLA-A*02:01-S976** |
|---|---|---|---|
| Temperature | 100K | 100K | 100K |
| Space group | $P\ 2_1\ 2_1\ 2_1$ | $P\ 2_1\ 2_1\ 2_1$ | $P\ 2_1\ 2_1\ 2_1$ |
| Cell Dimensions (a,b,c) (Å) | 60.16, 79.02, 110.56 | 60.47, 80.61, 111.33 | 60.34, 80.75, 110.71 |
| Resolution (Å) | 47.87 – 1.78 (1.82 – 1.78) | 78.37 – 1.79 (1.83 – 1.79) | 78.33 – 1.90 (1.95 – 1.90) |
| Total number of observations | 370593 (19069) | 353384 (20331) | 365840 (21514) |
| Number of unique observations | 50802 (2682) | 52085 (3041) | 42888 (2590) |
| Multiplicity | 7.3 (7.1) | 6.8 (6.7) | 8.5 (8.3) |
| Data completeness (%) | 99.6 (92.9) | 100 (100) | 99.3 (90.5) |
| $I/\sigma_I$ | 11.6 (1.8) | 11.3 (1.8) | 11.7 (1.8) |
| $R_{pim}{}^a$ (%) | 4.1 (43.5) | 3.7 (49.6) | 3.9 (45.7) |
| CC½ (%) | 99.8 (57.5) | 99.6 (66.9) | 99.7 (59.5) |
| **Refinement Statistics** | | | |
| Non-hydrogen atoms, Proteins | 2242 | 3331 | 3204 |
| Non-hydrogen atoms, Water | 406 | 255 | 356 |
| $R_{factor}{}^b$ (%) | 17.7 | 18.3 | 17. 1 |
| $R_{free}{}^b$ (%) | 20.7 | 21.9 | 20.6 |
| Rms deviations from ideality | | | |
| Bond lengths (Å) | 0.006 | 0.008 | 0.010 |
| Bond angles (°) | 0.828 | 0.975 | 1.000 |
| Ramachandran plot (%) | | | |
| Allowed region | 100 | 100 | 100 |
| Disallowed region | 0 | 0 | 0 |
| **PDB code** | 7T3G | 7T3W | 7SIS |

$^aR_{p.i.m} = \Sigma_{hkl}\,[1/(N-1)]^{1/2}\,\Sigma_i\,|\,I_{hkl,\,i} - \langle I_{hkl}\rangle\,|\,/\,\Sigma_{hkl}\,\langle I_{hkl}\rangle$. $^b\ R_{factor} = \Sigma_{hkl}\ ||\,F_o\,| - |\,F_c\,||\,/\,\Sigma_{hkl}|\,F_o\,|$ for all data except ≈ 5% which were used for $R_{free}$ calculation.

**Table S3. Thermal stability of the pHLA-A*02:01 complexes.**

| Peptide name | Tm (°C) ± S.E.M. |
|---|---|
| S367 | 46.05 ± 0.18 |
| S367-O | 57.55 ± 0.19 |
| S417 | 55.18 ± 0.67 |
| S417-O | 47.78 ± 0.70 |
| S976 | 50.50 ± 0.26 |
| S976-O | 50.10 ± 0.32 |

$T_m$ represents the mean thermal midpoint temperatures from two independent experiments (n=2). S.E.M represents the standard error of the mean from these data.

**References (29-37)**


29. E. J. Grant *et al.*, Broad CD8(+) T cell cross-recognition of distinct influenza A strains in humans. *Nat Commun* **9**, 5427 (2018).
30. C. L. Hsieh *et al.*, Structure-based design of prefusion-stabilized SARS-CoV-2 spikes. *Science* **369**, 1501-1505 (2020).
31. D. S. M. Chatzileontiadou, C. Szeto, D. Jayasinghe, S. Gras, Protein purification and crystallization of HLA-A *02:01 in complex with SARS-CoV-2 peptides. *STAR Protoc* **2**, 100635 (2021).
32. D. Aragao *et al.*, MX2: a high-flux undulator microfocus beamline serving both the chemical and macromolecular crystallography communities at the Australian Synchrotron. *J Synchrotron Radiat* **25**, 885-891 (2018).
33. W. Kabsch, Xds. *Acta Crystallogr D Biol Crystallogr* **66**, 125-132 (2010).
34. A. J. McCoy *et al.*, Phaser crystallographic software. *J Appl Crystallogr* **40**, 658-674 (2007).
35. The CCP4 suite: programs for protein crystallography. *Acta Crystallogr D Biol Crystallogr* **50**, 760-763 (1994).
36. P. Emsley, B. Lohkamp, W. G. Scott, K. Cowtan, Features and development of Coot. *Acta Crystallogr D Biol Crystallogr* **66**, 486-501 (2010).
37. B. E. Bricogne G., Brandl M., Flensburg C., Keller P., Paciorek W.,, S. A. Roversi P, Smart O.S., Vonrhein C., Womack T.O., Buster version 2.10. *Cambridge, United Kingdom: Global Phasing Ltd.*, (2011).
